# Protective Efficacy of Gastrointestinal SARS-CoV-2 Delivery Against Intranasal and Intratracheal SARS-CoV-2 Challenge in Rhesus Macaques

**DOI:** 10.1101/2021.09.13.460191

**Authors:** Jingyou Yu, Natalie D. Collins, Noe B. Mercado, Katherine McMahan, Abishek Chandrashekar, Jinyan Liu, Tochi Anioke, Aiquan Chang, Victoria M. Giffin, David L. Hope, Daniel Sellers, Felix Nampanya, Sarah Gardner, Julia Barrett, Huahua Wan, Jason Velasco, Elyse Teow, Anthony Cook, Alex Van Ry, Laurent Pessaint, Hanne Andersen, Mark G. Lewis, Christian Hofer, Donald S. Burke, Erica K. Barkei, Hannah A.D. King, Caroline Subra, Diane Bolton, Kayvon Modjarrad, Nelson L. Michael, Dan H. Barouch

**Author notes:** Co-First Authors (Jingyou Yu, Natalie D. Collins). Correspondence: Dan H. Barouch.

## Abstract

Live oral vaccines have been explored for their protective efficacy against respiratory viruses, particularly for adenovirus serotypes 4 and 7. The potential of a live oral vaccine against severe acute respiratory syndrome coronavirus 2 (SARS-CoV-2), however, remains unclear. In this study, we assessed the immunogenicity of live SARS-CoV-2 delivered to the gastrointestinal tract in rhesus macaques and its protective efficacy against intranasal and intratracheal SARS-CoV-2 challenge. Post-pyloric administration of SARS-CoV-2 by esophagogastroduodenoscopy resulted in limited virus replication in the gastrointestinal tract and minimal to no induction of mucosal antibody titers in rectal swabs, nasal swabs, and bronchoalveolar lavage. Low levels of serum neutralizing antibodies were induced and correlated with modestly diminished viral loads in nasal swabs and bronchoalveolar lavage following intranasal and intratracheal SARS-CoV-2 challenge. Overall, our data show that post-pyloric inoculation of live SARS-CoV-2 is weakly immunogenic and confers partial protection against respiratory SARS-CoV-2 challenge in rhesus macaques.

**Importance:** SARS-CoV-2 remains a global threat, despite the rapid deployment but limited coverage of multiple vaccines. Alternative vaccine strategies that have favorable manufacturing timelines, greater ease of distribution and improved coverage may offer significant public health benefits, especially in resource-limited settings. Live oral vaccines have the potential to address some of these limitations; however no studies have yet been conducted to assess the immunogenicity and protective efficacy of a live oral vaccine against SARS-CoV-2. Here we report that oral administration of live SARS-CoV-2 in non-human primates may offer prophylactic benefits, but that formulation and route of administration will require further optimization.

## Introduction

Coronavirus disease 2019 (COVID-19) has claimed millions of lives since its emergence in late 2019. Rapid and broad deployment of safe, effective and affordable vaccines will be the key to end the pandemic (*1, 2*). Multiple SARS-CoV-2 vaccines—including two mRNA vaccines and two adenoviral vectored vaccines—have advanced to emergency authorization or full approval at an unprecedented pace. Yet the wide gap in global availability of vaccines and the emergence of virus variants necessitate additional vaccine approaches (*1*).

Live oral vaccines have long been explored for their utility to curb infectious diseases. Immunologically, the gastrointestinal (GI) tract is one of the largest lymphoid organs in the body, comprised of organized lymphoid tissue and large populations of scattered innate and adaptive effector cells, including IgA-secreting plasma cells, CD4+ and CD8+ T cells, regulatory T cells, and γδ T cells (*3, 4*). Orally administered live vaccines may therefore elicit different immune responses than non-replicating gene-based vaccines, and the GI delivery route may be a means of attenuation (*5*). Direct administration of antigens at mucosal surfaces is an efficient approach to inducing a potent mucosal immune response (*6*). Logistically, live oral vaccines allow for simplified development, rapid production and distribution and ease of administration (*7*). Live virus production can be scaled up in cell culture systems without the need for complex inactivation and purification steps. Vaccination procedures are free of needles and there is often no need for specially trained medical personnel (*8*). Moreover, live oral vaccines are typically cost-effective. The replicating feature of live viruses can allow for administration of a lower dose to achieve immunity. As such, oral vaccines may be preferable in resource-limited settings.

To date, several human oral vaccines have been licensed that contain live viruses. The US Department of Defense (DoD) and National Institutes of Health (NIH) developed co-administered live oral vaccines against adenovirus serotypes 4 and 7 (Ad4 and Ad7) in the 1970s (*9, 10*) and again in 2011 when the vaccine was re-formulated (*11-13*). These two vaccines contain wild-type virus with an enteric coating to protect against degradation from the low pH of gastric acid as they pass through to the lower GI tract (*12, 14, 15*). GI administration of Ad4 and Ad7 attenuates the viruses and induces serum-neutralizing antibodies that protect against subsequent type-specific respiratory infection (*12, 14, 15*). Both vaccines have been shown to be safe, do not disseminate systemically—evident by absence of vaccine virus in blood or urine— and provide more than 90% efficacy over the course of 8 to 10 weeks (*11, 12, 14, 15*). Recent data have revealed that the Ad4/Ad7 live oral vaccine elicited immune responses are durable for at least 6 years (*16*). Oral vaccines have also been developed for GI viruses, such as rotavirus and poliovirus, which have been in use for decades in children and have consistently demonstrated high safety, immunogenicity, and efficacy profiles (*17-20*).

Given the success of the live oral Ad4 and Ad7 vaccines and the demonstration of the presence of the angiotensin-converting enzyme 2 (ACE2) receptor, the primary receptor for SARS-CoV-2, throughout the GI tract mucosa (*21*), we performed a proof-of-concept study to assess the immunogenicity and protective efficacy of GI delivery of live SARS-CoV-2 in rhesus macaques. Delivery of 1×10^6^ 50% tissue culture infectious dose (TCID50) virus to the duodenum by endoscopy caused a transient infection with localized replication in the GI tract and was associated with modest immunogenicity and partial protection against intranasal and intratracheal SARS-CoV-2 challenge.

## Results

### Limited SARS-CoV-2 Replication in the Gastrointestinal Tract

To determine the immunogenicity and protective efficacy of the GI delivery of SARS-CoV-2, we inoculated 21 rhesus macaques with 1×10^6^ 50% tissue culture infectious dose (TCID50) SARS-CoV-2 from the WA1/2020 strain (NR-52281; BEI Resources) (N=9) or PBS sham controls (N=12) by esophagogastroduodenoscopy (EGD). The virus inoculum was 2 ml of live virus in PBS and was delivered to the proximal duodenum on day 0.

Viral shedding was quantified on study days 1, 2, 4, 7, 14, 21 and 28 by genomic (gRNA) or envelope (E) subgenomic (sgRNA) RT-PCR assays (*22*). Viral shedding in the stool was observed in 7 out of 9 vaccinated macaques by gRNA assays on day 1 post-inoculation, but only one macaque had sustained viral shedding in stool for more than 21 days (Fig. 1A). Additionally, virus was observed by gRNA assays from rectal swabs (RS) in 4 out of 9 macaques, with detectable virus in two macaques at 21 days post-immunization (Fig. 1B). In contrast, virus was not detected in sham control macaques (Fig. 1A and 1B). Similar but limited viral shedding was observed by sgRNA assays in the vaccinated animals but not the sham controls (Fig. 1C and 1D). However, we did not observe virus in serum, saliva, bronchoalveolar lavage (BAL) or nasal swabs (NS) (data not shown). On day 1, vaccinated animals had a median 3.49 log_10_ viral copies per gram stool, whereas the sham animals had no detectable virus (P<0.00001, two-sided Mann-Whitney tests) (Fig. 1E). These data suggest that the virus inoculum was rapidly excreted with limited virus replication in the GI tract.

**Figure 1.**
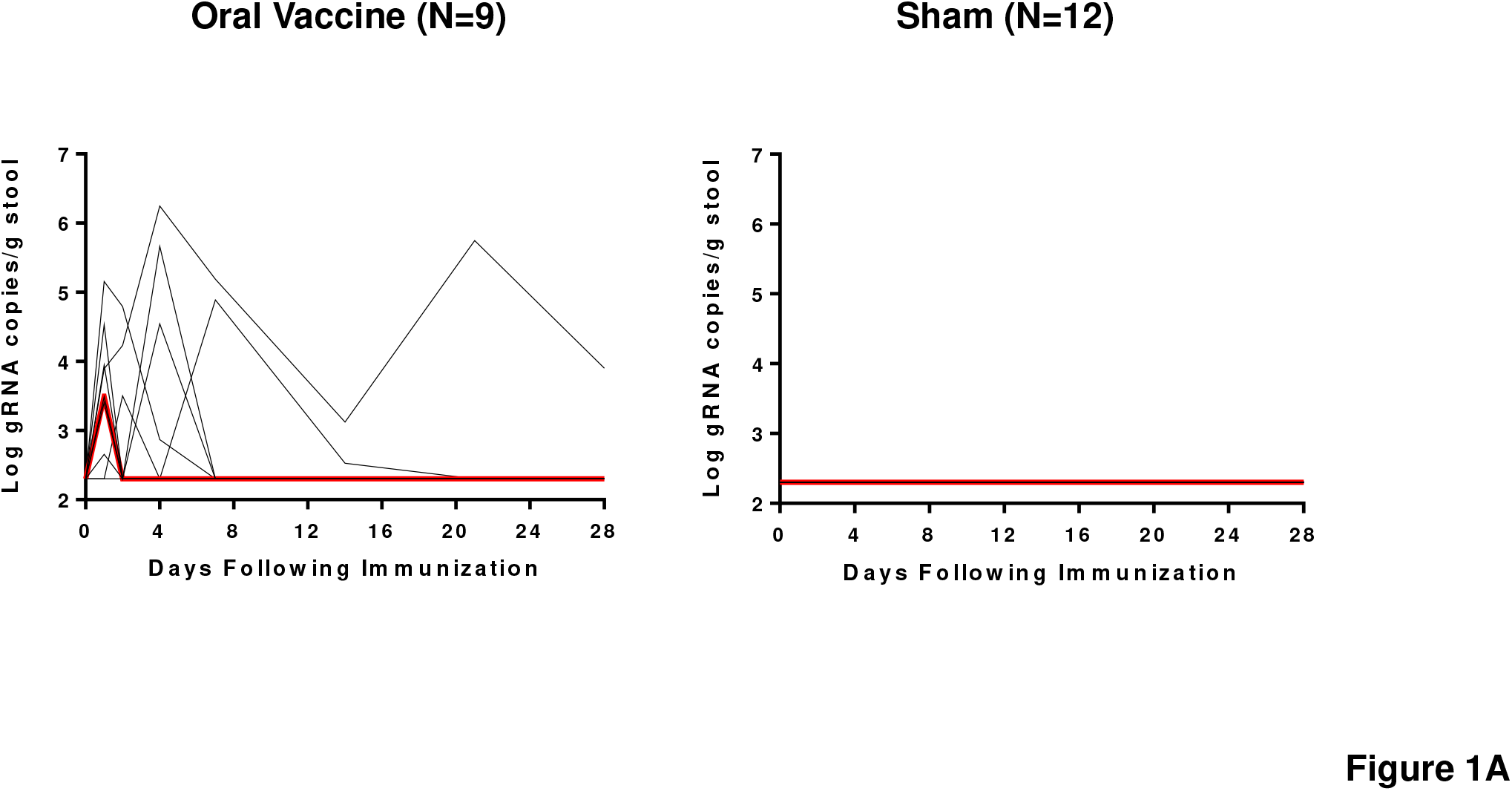
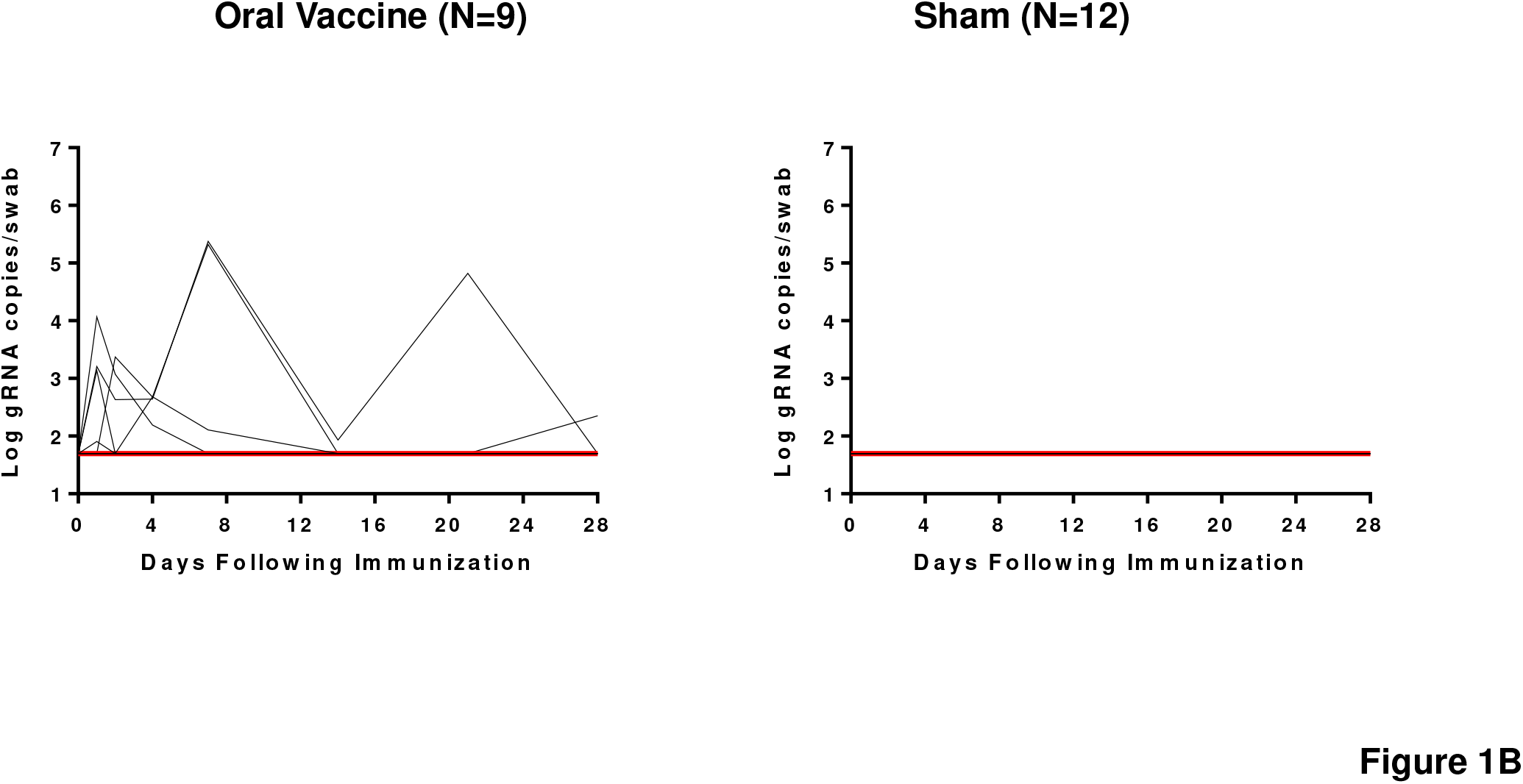
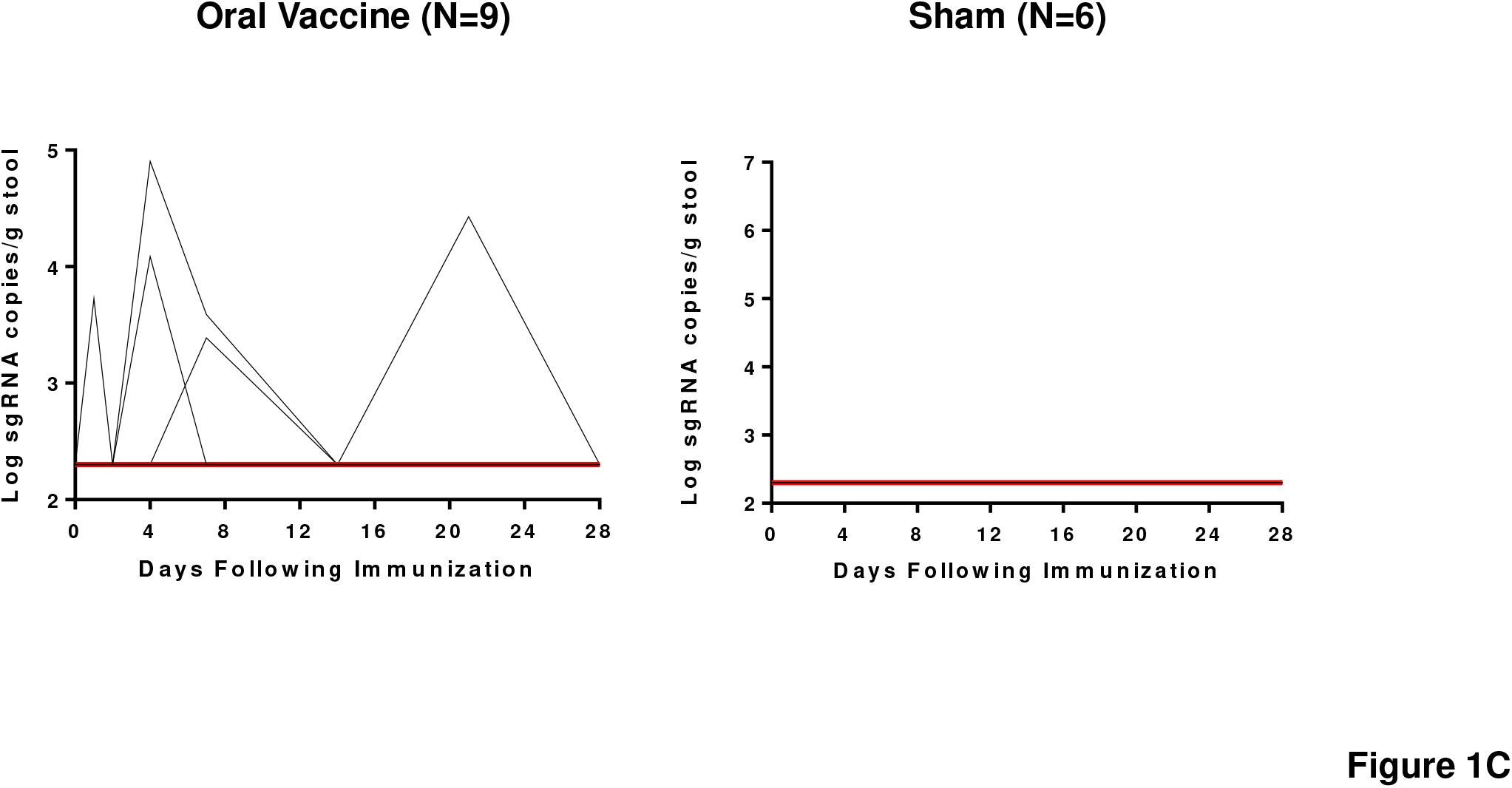
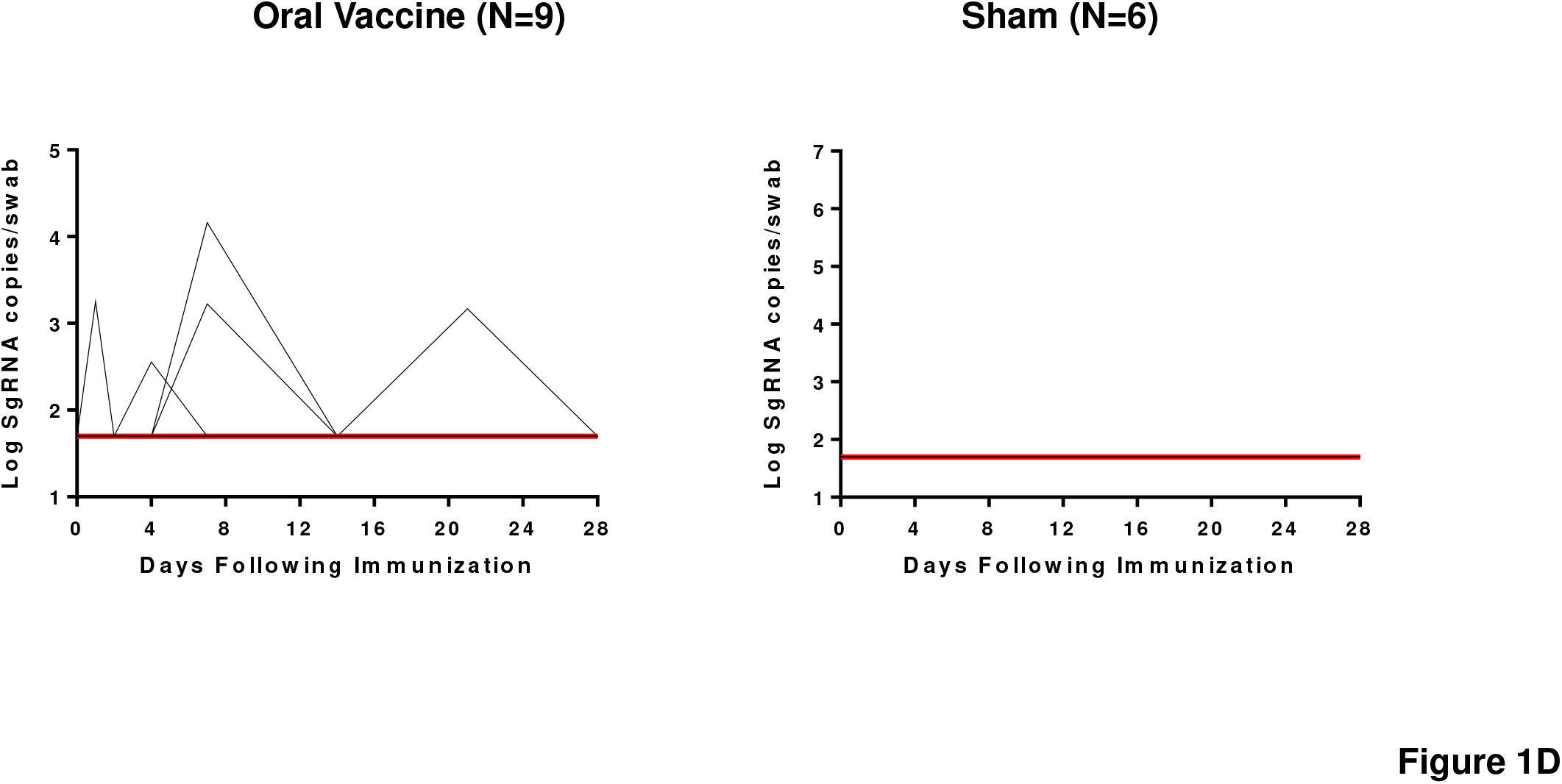
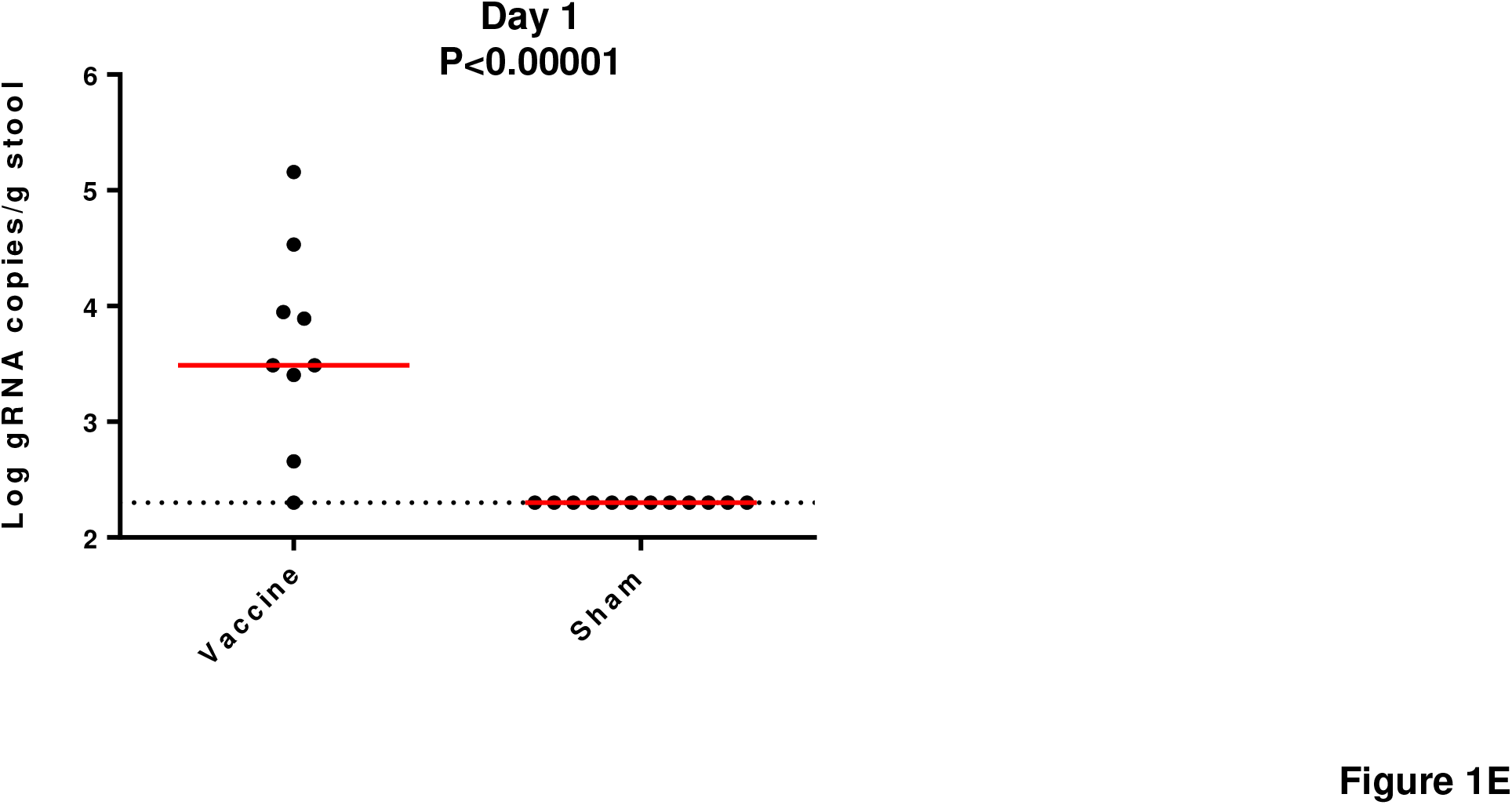
Viral shedding in rhesus macaques following live vaccine EGD administration. Rhesus macaques were administered 10^6^ TCID50 SARS-CoV-2GI via EGD. (A) Log10 gRNA copies/g stool (limit 200 copies/ml) or (C) Log10 sgRNA copies/g stool were assessed in stools in sham controls and in vaccinated animals following challenge. (B) Log10 gRNA copies/swab or (D) Log10 sgRNA copies/swab (limit 50 copies/swab) were assessed in rectal swabs (RS) in sham controls and in vaccinated animals following challenge. Red lines reflect median values. (E) Peak viral loads in stool on day 1 following vaccination. Red lines reflect median viral loads. P-values indicate two-sided Mann-Whitney tests.

### Immunogenicity of GI Delivery of SARS-CoV-2 Live Vaccine in Rhesus Macaques

Four weeks after vaccination, we observed low serum pseudovirus neutralizing antibody (NAb) titers in 7 of 9 vaccinated macaques (Fig. 2), whereas the sham animals had undetectable NAb titers. NAb titers in mucosal specimens, including NS, BAL, RS, and stool, were below the limit of detection (data not shown). We assessed T cell responses in peripheral blood mononuclear cells (PBMCs) at week 4 post-inoculation and found undetectable responses to pooled S peptides in both vaccinated and unvaccinated animals by IFN-γ ELISPOT assays and intracellular cytokine staining (ICS) assays (data not shown). Together, these data suggest that the GI delivery of SARS-CoV-2 generated modest levels of serum neutralizing antibodies but undetectable mucosal immune responses and cellular immune responses.

**Figure 2.**
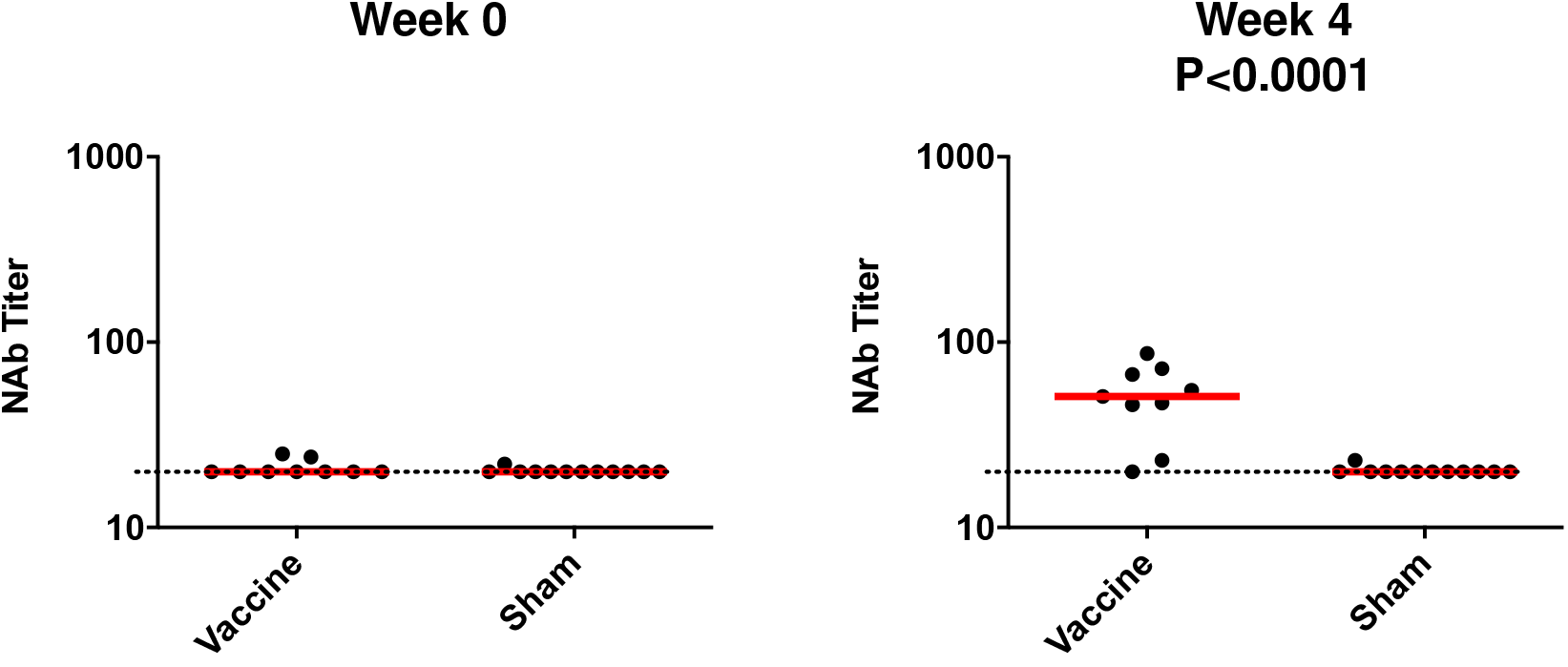
Humoral immune responses in vaccinated rhesus macaques. Humoral immune responses were assessed at weeks 0 and 4 by pseudovirus neutralization assays. Red bars reflect median responses. Dotted lines reflect assay limit of detection.

### Protective Efficacy Against SARS-CoV-2 Challenge

At week 4 post-inoculation, all animals were challenged with 10^5^ TCID50 of SARS-CoV-2 WA1/2020, administered in a 2 ml volume by the intranasal (IN) and intratracheal (IT) routes. Following challenge, we assessed viral loads in the BAL and NS (*22, 23*). High levels of sgRNA were observed in the sham controls with a median peak of 4.79 (range 2.61-5.69) log_10_ sgRNA copies/ml in BAL and a median peak of 6.21 (range 3.30-6.82) log_10_ sgRNA copies/swab in NS (Fig. 3A and 3B). Lower viral loads were observed in the vaccinated macaques (Fig. 3A and 3B), with 1.61 and 1.59 log_10_ reductions of median peak sgRNA in BAL and NS, respectively (P=0.0040 and P=0.0093, two-sided Mann-Whitney tests) (Fig. 3C). These data demonstrate that the GI delivered SARS-CoV-2 provided partial but modest protection against respiratory SARS-CoV-2 challenge.

**Figure 3.**
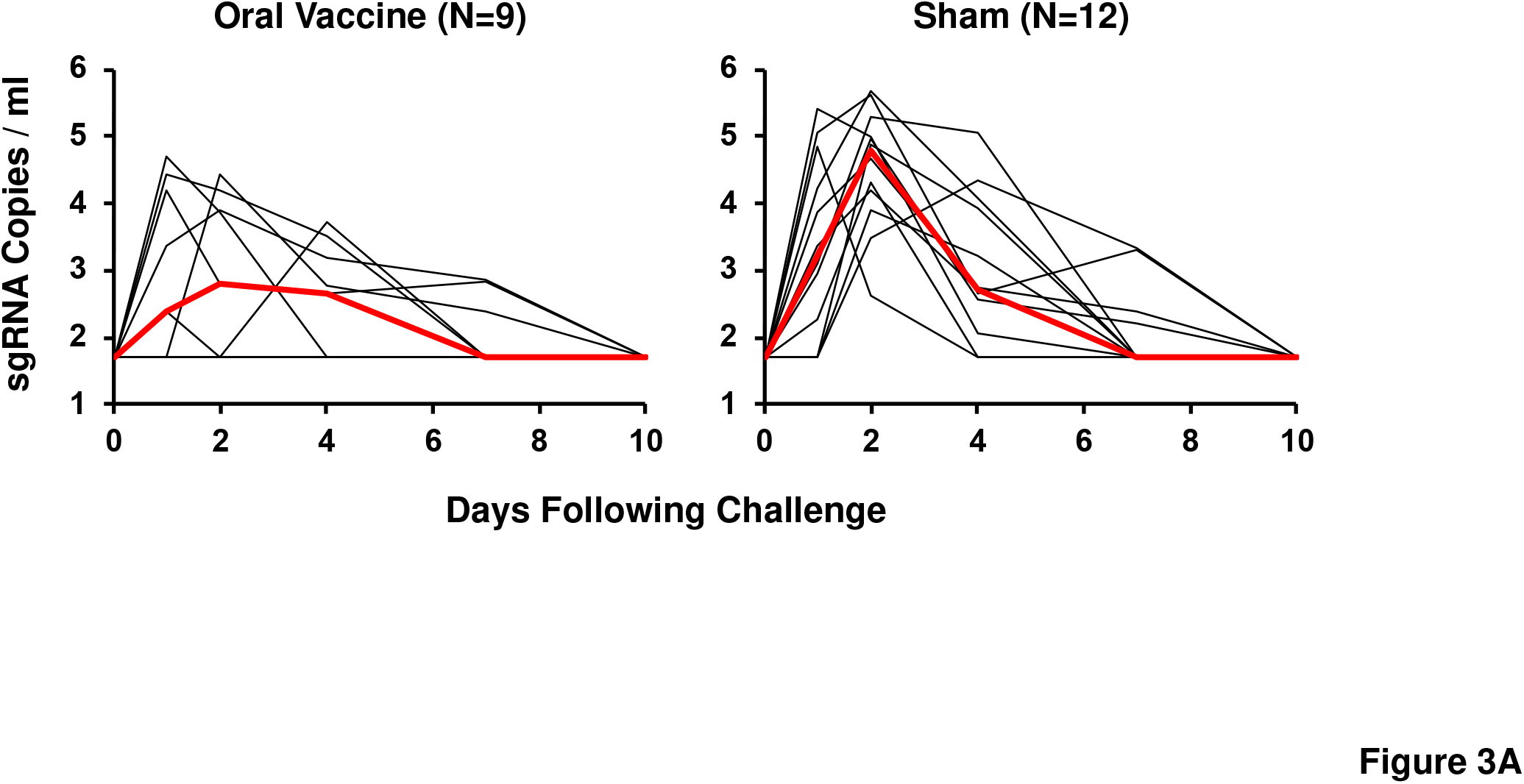
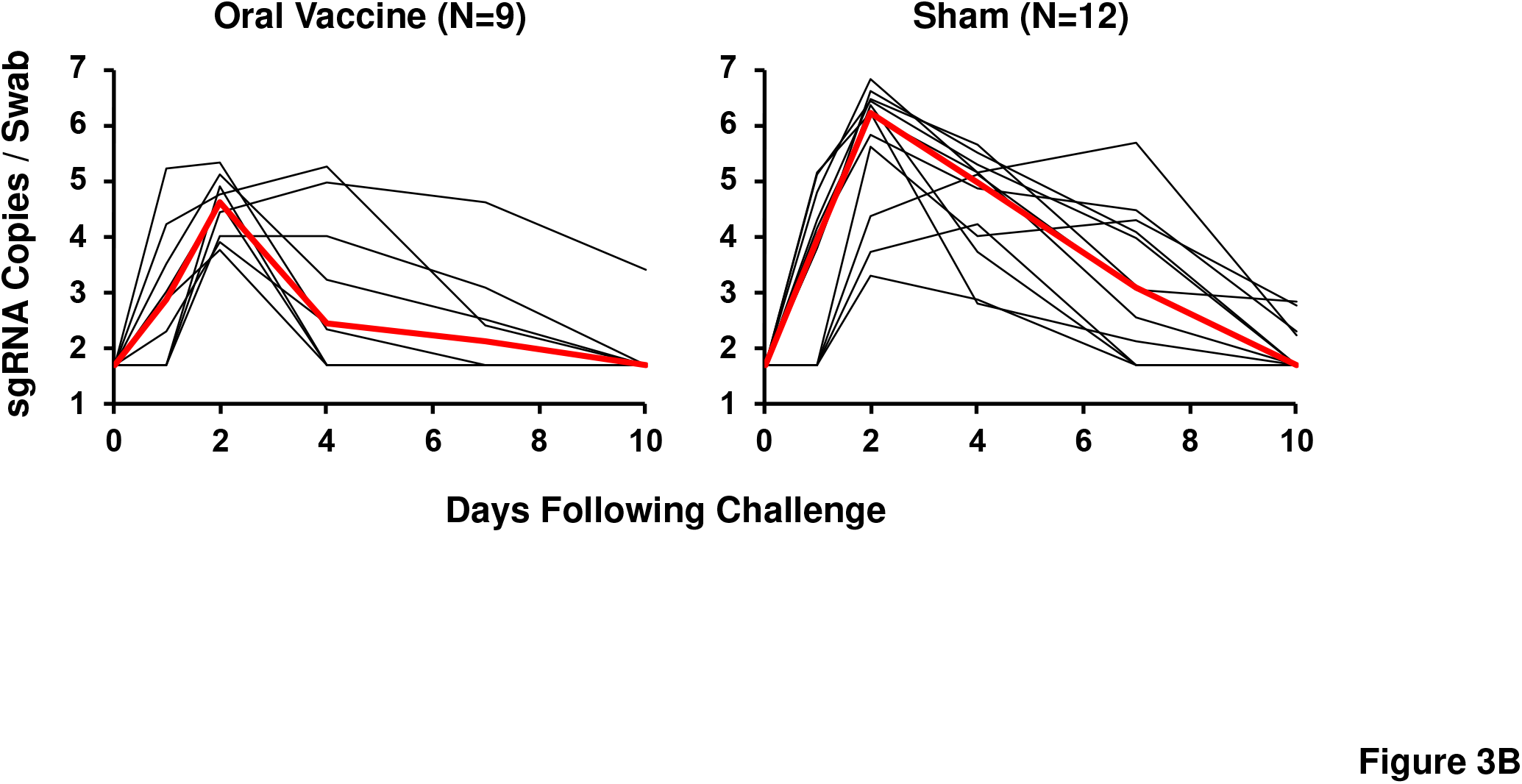
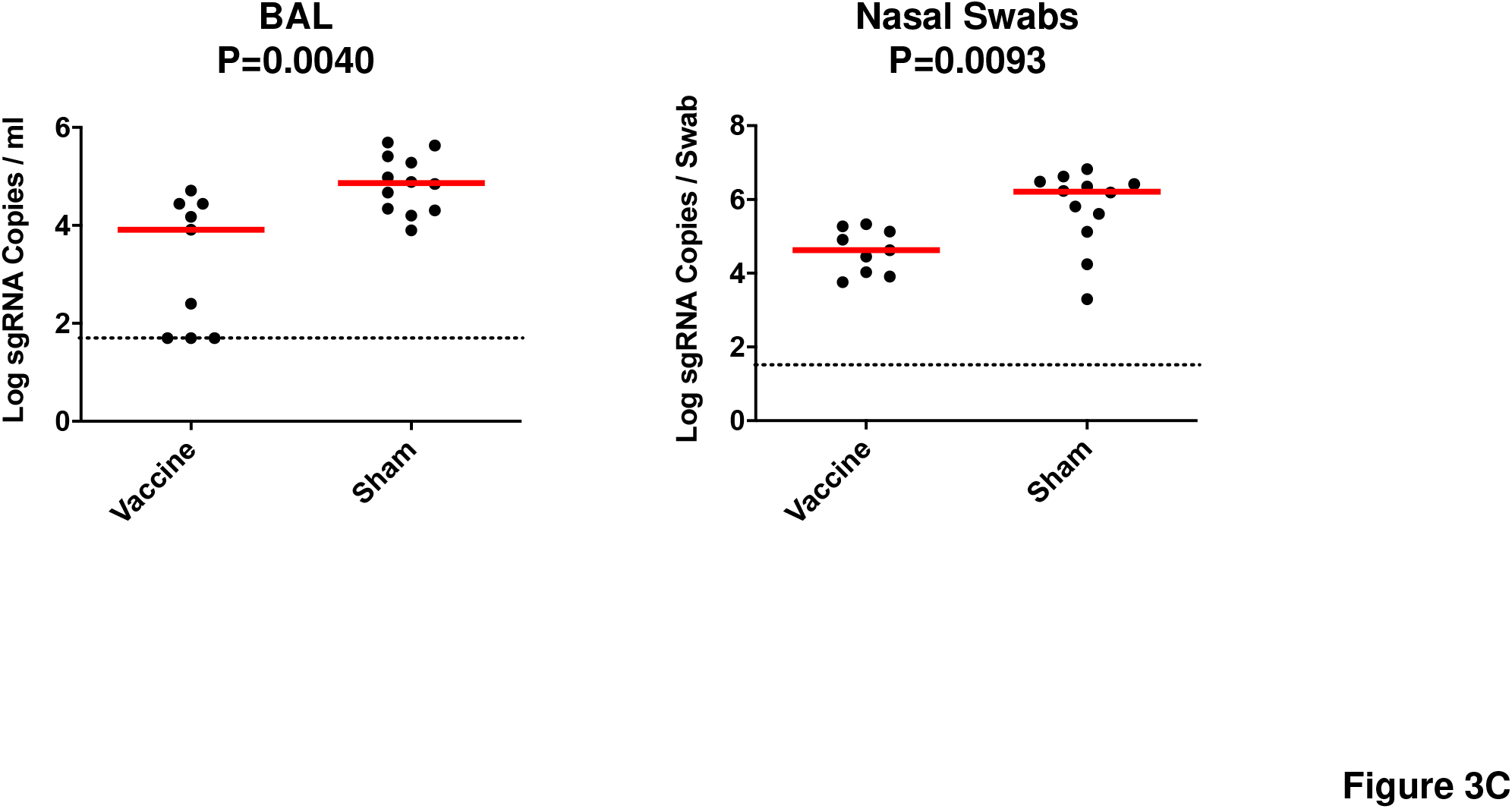
Viral loads in rhesus macaques following SARS-CoV-2 challenge. Rhesus macaques were challenged by the intranasal and intratracheal route with 10^5^ TCID50 SARS-CoV-2. (A) Log10 sgRNA copies/ml (limit 50 copies/ml) were assessed in bronchoalveolar lavage (BAL) in sham controls and in vaccinated animals following challenge. (B) Log10 sgRNA copies/swab (limit 50 copies/swab) were assessed in nasal swabs (NS) in sham controls and in vaccinated animals following challenge. Red lines reflect median values. (C) Peak viral loads in BAL and NS following challenge. Peak viral loads occurred on day 2 following challenge. Red lines reflect median viral loads. P-values indicate two-sided Mann-Whitney tests.

On day 14 following challenge, histopathology revealed minimal to mild interstitial pneumonia in all animal groups, characterized by type II pneumocyte hyperplasia, perivascular inflammation and/or vasculitis of small to medium-sized vessels, and thickening of alveolar septae by fibrin and/or mononuclear inflammatory cells (Fig. 4). No clear difference in pulmonary pathology was noted between the vaccinated aniamals and sham controls.

**Figure 4.**
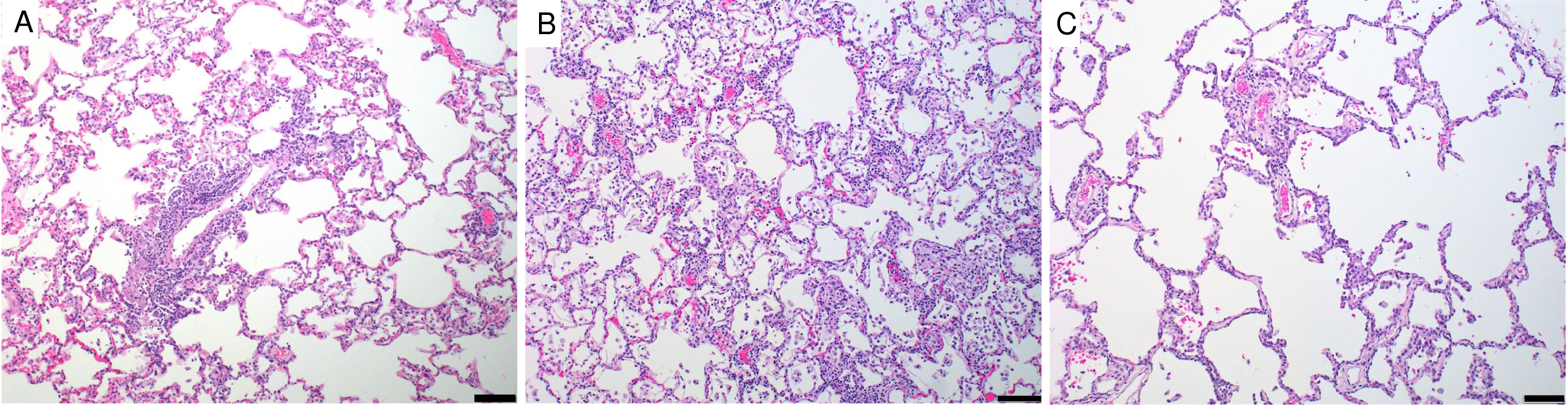
Histopathologic examination following SARS-CoV-2 challenge. Lung tissues were collected at necropsy on day 14 post-challenge, fixed with neutral buffered formalin, and stained with hematoxylin and eosin (H&E) for standard microscopic examination. Representative lung tissue sections from the PBS control (A), high-dose (10^6^ TCID50) vaccinated (B) low-dose (10^4^ TCID50) vaccinated (C) SARS-CoV-2 challenged rhesus macaques. Minimal to mild interstitial pneumonia is characterized by inflammatory cellular infiltrates and type II pneumocyte hyperplasia. Scale bars: 100 µm.

### Immune Correlates of Protection

Given the observed protection, we assessed immune correlates of protection. As shown in Fig. 5A, the log_10_ pseudovirus NAb titer at week 4 inversely correlated with peak log_10_ sgRNA copies/ml in both BAL (R=-0.6165, P=0.0029) and NS (R=-0.3693, P=0.0994) (Fig. 5A). The less robust correlation with viral loads in NS compared with viral loads in BAL is consistent with prior studies (*24, 25*). As shown in Fig. 5B, peak viral shedding in stool did not correlate with peak log_10_ sgRNA copies/ml in BAL and NS.

**Figure 5.**
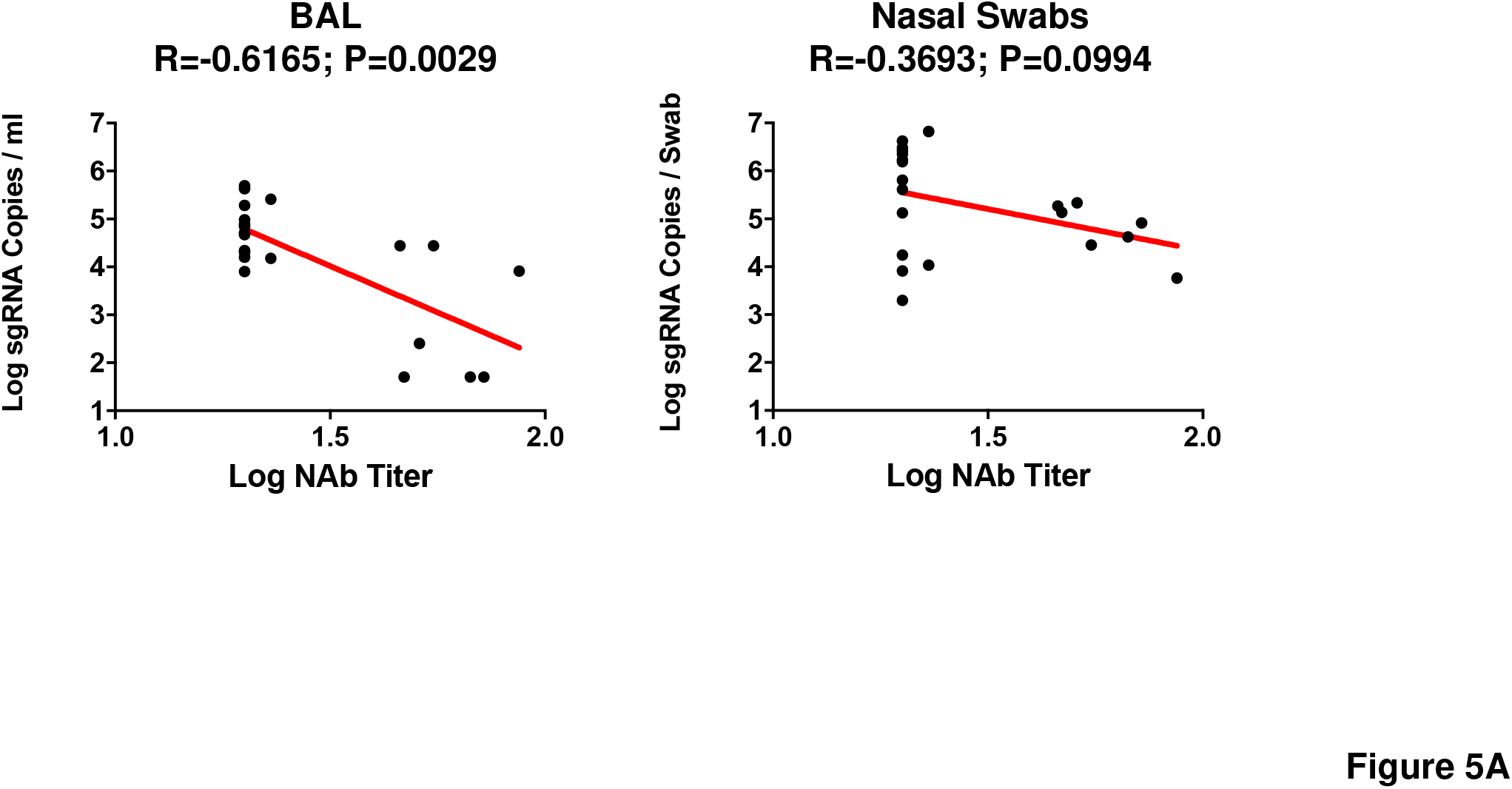
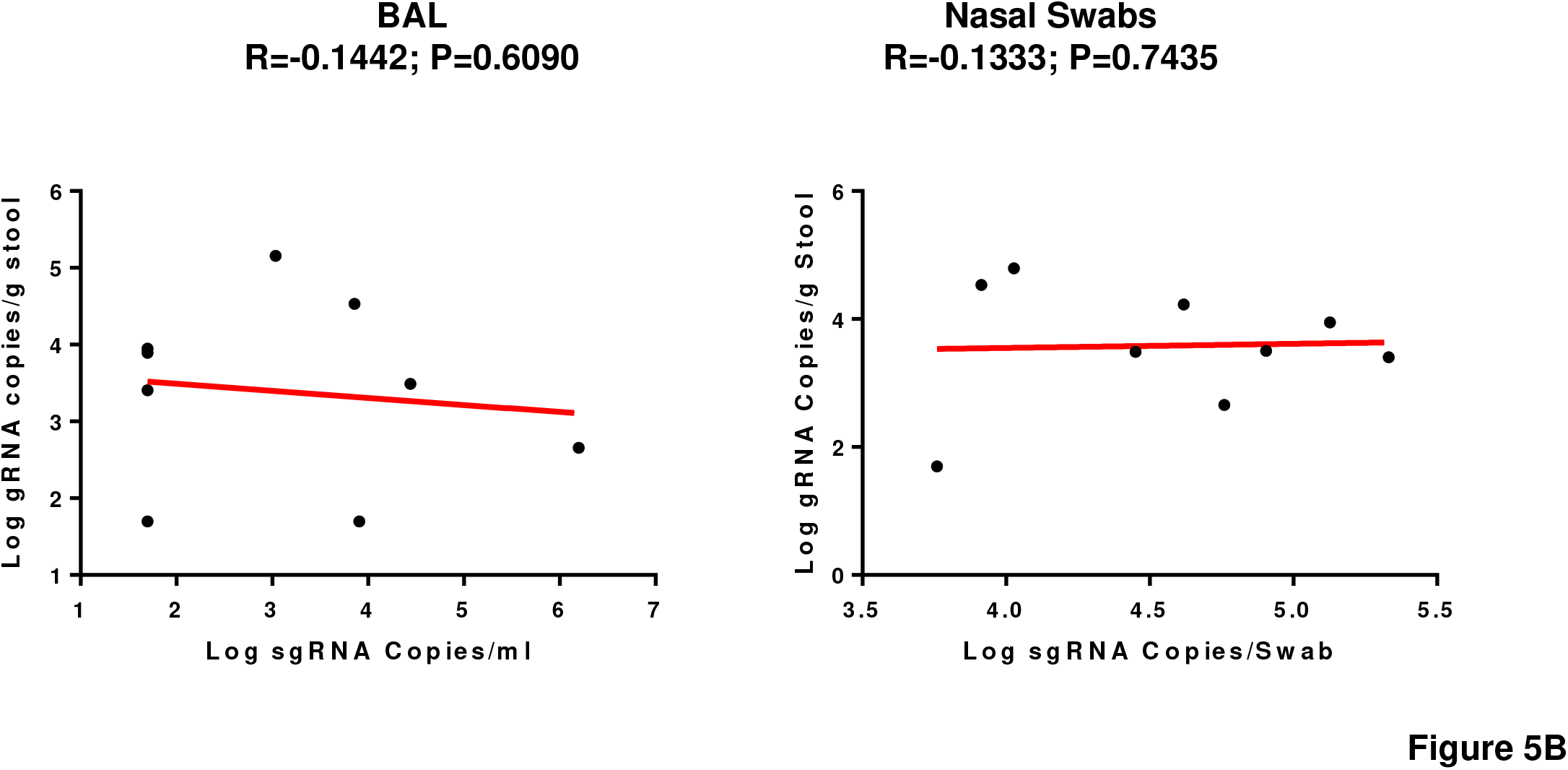
Immune correlates of protection. (A) Correlations of pseudovirus NAb titers at week 4 with log peak sgRNA copies/ml in BAL and NS following challenge. (B) Correlations of log peak sgRNA copies/ml in BAL and NS with log peak gRNA copies/g stool. Red lines reflect the best-fit relationship between these variables. P and R values reflect two-sided Spearman rank-correlation tests.

## Discussion

In this study, we demonstrate that GI delivery of live 1×10^6^ TCID50 SARS-CoV-2 elicited modest immune responses and provided partial protection against intranasal and intratracheal challenge with SARS-CoV-2. Moreover, serum neutralizing antibody titers correlated with protective efficacy. These data provide proof-of-concept that an orally administered vaccine can protect against respiratory SARS-CoV-2 challenge, but the limited immunogenicity and protective efficacy observed here suggests that the oral vaccine approach will require optimization.

SARS-CoV-2 has been shown to productively infect human and macaque GI tract (*26-28*), specifically enterocytes (*29-31*), and infection is frequently associated with clinical symptoms in humans (*32*). We thus hypothesized that the replication of the live viral vaccine in the gut may lead to induction of systemic and mucosal immunity. We observed rapid excretion of the virus with minimal replication in the GI tract, which may explain the poor immunogenicity and limited protection. In contrast, Jiao et al. recently reported that intragastric inoculation of 1 × 10^7^ PFU SARS-CoV-2 in rhesus macaques resulted in a productive and sustained viral infection in GI tract (*28*). This could reflect different inoculum doses, administration techniques, or animal cohorts.

Data on expression of ACE2 receptor in the stomach and GI tract is limited. Available data suggest abundant expression in the small intestines (*21*). Taken together with minimal replication of SARS-CoV2 in the GI tract, but clear correlation of serum NAb with protection in the BAL and NS, it is likely that the limited immune responses were due to an inadequate antigenic load in the GI tract. Therefore, optimization of the oral vaccine formulation, including the use of encapsulation and buffers for improved controlled delivery of SARS-CoV-2 to the GI tract, with adequate time for viral replication in a hospitable micro-environment, may allow more effective delivery of an oral vaccine. Higher doses or repetitive doses may also prove useful.

In summary, our data show that a single post-pyloric administration of live SARS-CoV-2 by EGD elicited detectable serum NAb titers and partially protected against respiratory SARS-CoV-2 challenge in rhesus macaques. Optimization of the current strategy, with encapsulation and extended delivery systems, as well as improvements in dosage and schedule will be required for a live, oral, SARS-CoV2 vaccine.

## Acknowledgements

We thank Robert Kushner, Daniel Sellers, Owen Sanborn, Kunza Ahmad, for generous advice, assistance, and reagents. This project was funded by the Henry Jackson Foundation (2018-CHEDA-001-948100), the Musk Foundation, and the Ragon Institute of MGH, MIT, and Harvard.

## Author Contributions

D.H.B., J.Y., N.D.C. K.M., N.L.M., D.L.B. designed the study and reviewed all data. J.Y., N.B.M., K.M., J.L., A.C., J.L., A.C., D.L.H., V.M.G., F.N., S.P., H.W, C.S., H.A.D.K. performed the immunologic and virologic assays. E.K.B performed histological studies. J.V., E.T., A.C., A.V.R., L.P., H.A., and M.G.L. led the clinical care of the animals. J.Y., N.D.C., and D.H.B. wrote the paper with all co-authors.

## Material and Methods

### Animals, virus stocks, and study design

21 outbred Indian-origin adult male and female rhesus macaques (*Macaca mulatta*) ages 6-14 years old were randomly allocated to groups. All animals were housed at Bioqual, Inc. (Rockville, MD). Animals were EGD administered into duodenum with 1 x10^6^ TCID50 SARS-CoV-2 and then challenged with 10^5^ TCID50 of WA1/2020 on day 28. The WA1/2020 (USA-WA1/2020; BEI Resources; NR-5228) challenge stock was grown in VeroE6 cells and deep sequenced as described previously (*33*). Deep sequencing of these stocks revealed no mutations in the Spike protein greater than >2.5% frequency. At the time of challenge, virus was administered as 1 ml by the intranasal (IN) route (0.5 ml in each nare) and 1 ml by the intratracheal (IT) route. All immunologic and virologic studies were performed blinded. Animal studies were conducted in compliance with all relevant local, state, and federal regulations and were approved by the Bioqual Institutional Animal Care and Use Committee (IACUC).

### EGD administration

The scope was slowly and trans-orally inserted, under direct vision. Once the endoscope was in the stomach, insufflation, aspiration, and suctioning were used to aid in finding the specified gastro-intestinal region (pyloric region, the duodenum, or the jejunum). Once the duodenum was identified, inoculum was administered through the instrument channel inlet. The channel was then flushed with 1-2 mL of sterile water. The endoscope was removed and cleaned in between animals with appropriate disinfectant. A new endoscope was used on another animal while the previous endoscope was disinfected.

### Pseudovirus-based virus neutralization assay

The SARS-CoV-2 pseudoviruses expressing a luciferase reporter gene were generated essentially as described previously (*24, 25, 33, 34*). Briefly, the packaging plasmid psPAX2 (AIDS Resource and Reagent Program), luciferase reporter plasmid pLenti-CMV Puro-Luc (Addgene), and spike protein expressing pcDNA3.1-SARS CoV-2 SΔCT of variants were co-transfected into HEK293T cells by lipofectamine 2000 (ThermoFisher). Pseudoviruses of SARS-CoV-2 variants were generated by using Wuhan/WIV04/2019strain (GISAID accession ID: EPI_ISL_402124). The supernatants containing the pseudotype viruses were collected 48 h post-transfection, which were purified by centrifugation and filtration with 0.45 µm filter. To determine the neutralization activity of the plasma or serum samples from participants, HEK293T-hACE2 cells were seeded in 96-well tissue culture plates at a density of 1.75 × 10^4^ cells/well overnight. Three-fold serial dilutions of heat inactivated serum or nasal swab, BAL, rectal swab or stools were prepared and mixed with 50 µL of pseudovirus. The mixture was incubated at 37°C for 1 h before adding to HEK293T-hACE2 cells. 48 h after infection, cells were lysed in Steady-Glo Luciferase Assay (Promega) according to the manufacturer’s instructions. SARS-CoV-2 neutralization titers were defined as the sample dilution at which a 50% reduction in relative light unit (RLU) was observed relative to the average of the virus control wells.

### ELISA

WA1/2020 RBD-specific binding antibodies were assessed by ELISA essentially as described previously (*24, 25, 33*). Briefly, 96-well plates were coated with 1µg/ml RBD protein (source: Aaron Schmidt) in 1X DPBS and incubated at 4°C overnight. After incubation, plates were washed once with wash buffer (0.05% Tween 20 in 1 X DPBS) and blocked with 350 µL Casein block/well for 2-3 h at room temperature. After incubation, block solution was discarded, and plates were blotted dry. Serial dilutions of heat-inactivated serum diluted in casein block were added to wells and plates were incubated for 1 h at room temperature, prior to three further washes and a 1 h incubation with a 1µg/ml dilution of anti-macaque IgG HRP (Nonhuman Primate Reagent Resource) or a 1:1000 dilution of anti-monkey IgA HRP (Novus) at room temperature in the dark. Plates were then washed three times, and 100 µL of SeraCare KPL TMB SureBlue Start solution was added to each well; plate development was halted by the addition of 100 µL SeraCare KPL TMB Stop solution per well. The absorbance at 450nm was recorded using a VersaMax microplate reader. For each sample, ELISA endpoint titer was calculated in Graphpad Prism software, using a four-parameter logistic curve fit to calculate the reciprocal serum dilution that yields an absorbance value of 0.2 at 450nm. Log10 endpoint titers are reported.

### IFN-γ enzyme-linked immunospot (ELISPOT) assay

ELISPOT assays were performed essentially as described previously (*24, 25, 33*). ELISPOT plates were coated with mouse anti-human IFN-γ monoclonal antibody from BD Pharmigen at 5 µg/well and incubated overnight at 4°C. Plates were washed with DPBS wash buffer (DPBS with 0.25% Tween20), and blocked with R10 media (RPMI with 10% heat inactivated FBS with 1% of 100x penicillin-streptomycin) for 1-4 h at 37°C. SARS-CoV-2 peptides pools from JPT were prepared & plated at a concentration of 1 µg/well, and 200,000 cells/well were added to the plate. The peptides and cells were incubated for 18-24 h at 37°C. All steps following this incubation were performed at room temperature. The plates were washed with ELISPOT wash buffer (11% 10x DPBS and 0.3% Tween20 in 1L MilliQ water) and incubated for 2 h with Rabbit polyclonal anti-human IFN-γ Biotin from U-Cytech (1 µg/mL). The plates were washed a second time and incubated for 2 h with Streptavidin-alkaline phosphatase from Southern Biotech (2 µg/mL). The final wash was followed by the addition of Nitro-blue Tetrazolium Chloride/5-bromo-4-chloro 3 ‘indolyphosphate p-toludine salt (NBT/BCIP chromagen) substrate solution for 7 min. The chromagen was discarded and the plates were washed with water and dried in a dim place for 24 h. Plates were scanned and counted on a Cellular Technologies Limited Immunospot Analyzer.

### Intracellular cytokine staining (ICS) assay

Multiparameter ICS assays were performed utilizing modification of described previously protocols (*24, 25, 33*).

### Genomic and Subgenomic RNA assay

SARS-CoV-2 E gene subgenomic RNA (sgRNA) and N gene genomic RNA (gRNA) were assessed by RT-PCR using primers and probes as previously described (*35, 36*). A standard was generated by first synthesizing a gene fragment of the subgenomic E gene (*36*). The gene fragment was subsequently cloned into a pcDNA3.1+ expression plasmid using restriction site cloning (Integrated DNA Technologies). The insert was in vitro transcribed to RNA using the AmpliCap-Max T7 High Yield Message Maker Kit (CellScript). Log dilutions of the standard were prepared for RT-PCR assays ranging from 1×1010 copies to 1×10-1 copies. Viral loads were quantified from bronchoalveolar lavage (BAL) fluid, nasal swabs (NS), rectal swabs (RS) and stool. RNA extraction was performed on a QIAcube HT using the IndiSpin QIAcube HT Pathogen Kit according to manufacturer’s specifications (Qiagen). The standard dilutions and extracted RNA samples were reverse transcribed using SuperScript VILO Master Mix (Invitrogen) following the cycling conditions described by the manufacturer, 25°C for 10 Minutes, 42°C for 1 Hour then 85°C for 5 Minutes. A Taqman custom gene expression assay (Thermo Fisher Scientific) was designed using the sequences targeting the E gene sgRNA (*36*). The sequences for the custom assay were as follows, sgLeadCoV2.Fwd: CGATCTCTTGTAGATCTGTTCTC, E_Sarbeco_R: ATATTGCAGCAGTACGCACACA, E_Sarbeco_P1 (probe): VIC-ACACTAGCCATCCTTACTGCGCTTCG-MGBNFQ. SARS-CoV-2 genomic RNA (gRNA) was targeted using N gene primers and probe, 2019-nCoV_N1-F: GACCCCAAAATCAGCGAAAT, 2019-nCoV_N1-R: TCTGGTTACTGCCAGTTGAATCTG, and 2019-nCoV_N1-P: FAM-ACCCCGCATTACGTTTGGTGGACC-BHQ1. Reactions were carried out in duplicate for samples and standards on the QuantStudio 6 and 7 Flex Real-Time PCR Systems (Applied Biosystems) with the thermal cycling conditions, initial denaturation at 95°C for 20 seconds, then 45 cycles of 95°C for 1 second and 60°C for 20 seconds. Standard curves were used to calculate genomic and subgenomic RNA copies per ml or per swab; the quantitative assay sensitivity was 50 copies per ml or per swab for both genomic and subgenomic assays. Sensitivity of the stool analysis was determined as 200 copies/ gram of stool.

### Histopathology

Necropsies were performed according to IACUC approved protocols at 14 days post infection. Lungs were perfused with 10% neutral-buffered formalin. Three tissue sections each from the right and left lung lobes were used to evaluate the lung pathology. Sections were processed routinely into paraffin wax, then sectioned at 5 µm, and resulting slides were stained with hematoxylin and eosin. All tissue slides were evaluated by a board-certified veterinary anatomic pathologist blinded to study group allocations.

### Statistical analyses

Comparisons of virologic and immunologic data was performed using GraphPad Prism 8.4.2 (GraphPad Software). Comparison of data between groups was performed using two-sided Wilcoxon rank-sum tests. Correlation analyses were performed either using two-sided Spearman rank-correlation tests or linear regression. P-values of less than 0.05 were considered significant.

